# The promiscuous estrogen receptor: evolution of physiological estrogens and response to phytochemicals and endocrine disruptors

**DOI:** 10.1101/228064

**Authors:** Michael E. Baker, Richard Lathe

## Abstract

Many actions of estradiol (E2), the principal physiological estrogen in vertebrates, are mediated by estrogen receptor-α (ERα) and ERβ. An important physiological feature of vertebrate ERs is their promiscuous response to several physiological steroids, including estradiol (E2), Δ^5^-androstenediol, 5α-androstanediol, and 27-hydroxycholesterol. A novel structural characteristic of Δ^5^-androstenediol, 5α-androstanediol, and 27-hydroxycholesterol is the presence of a C19 methyl group, which precludes the presence of an aromatic A ring with a C3 phenolic group that is a defining property of E2. The structural diversity of these estrogens can explain the response of the ER to synthetic chemicals such as bisphenol A and DDT, which disrupt estrogen physiology in vertebrates, and the estrogenic activity of a variety of plant-derived chemicals such as genistein, coumestrol, and resveratrol. Diversity in the A ring of physiological estrogens also expands potential structures of industrial chemicals that can act as endocrine disruptors. Compared to E2, synthesis of 27-hydroxycholesterol and Δ^5^-androstenediol is simpler, leading us, based on parsimony, to propose that one or both of these steroids or a related metabolite was a physiological estrogen early in the evolution of the ER, with E2 assuming this role later as the canonical estrogen. In addition to the well-studied role of the ER in reproductive physiology, the ER also is an important transcription factor in non-reproductive tissues such as the cardiovascular system, kidney, bone, and brain. Some of these ER actions in non-reproductive tissues appeared early in vertebrate evolution, long before mammals evolved.

## 1. Introduction

### 1.1 Promiscuity in steroids that activate the estrogen receptor

As our understanding of ligand-protein interactions has expanded, appreciation for the role of promiscuity as a force in evolution has increased [1-5]. The term ‘promiscuity’ is akin to that of ‘biological messiness’ [6]. We use promiscuity here as the capacity of the estrogen receptor (ER) to bind physiological steroids with diverse structures. In particular, the A ring of estradiol (E2), the main physiological estrogen, differs from the A ring in Δ^5^-androstenediol, 5α-androstanediol, and 27-hydroxycholesterol (Figure 1), as well as from the A ring of other classes of vertebrate adrenal and sex steroids (Figure 2). Indeed, the aromatic A ring and C3 phenolic group, characteristic of E2, have been exploited to synthesize ‘synthetic estrogens’ that are used to treat estrogen-dependent diseases (Figure 3) [7, 8].

**Figure 1.**
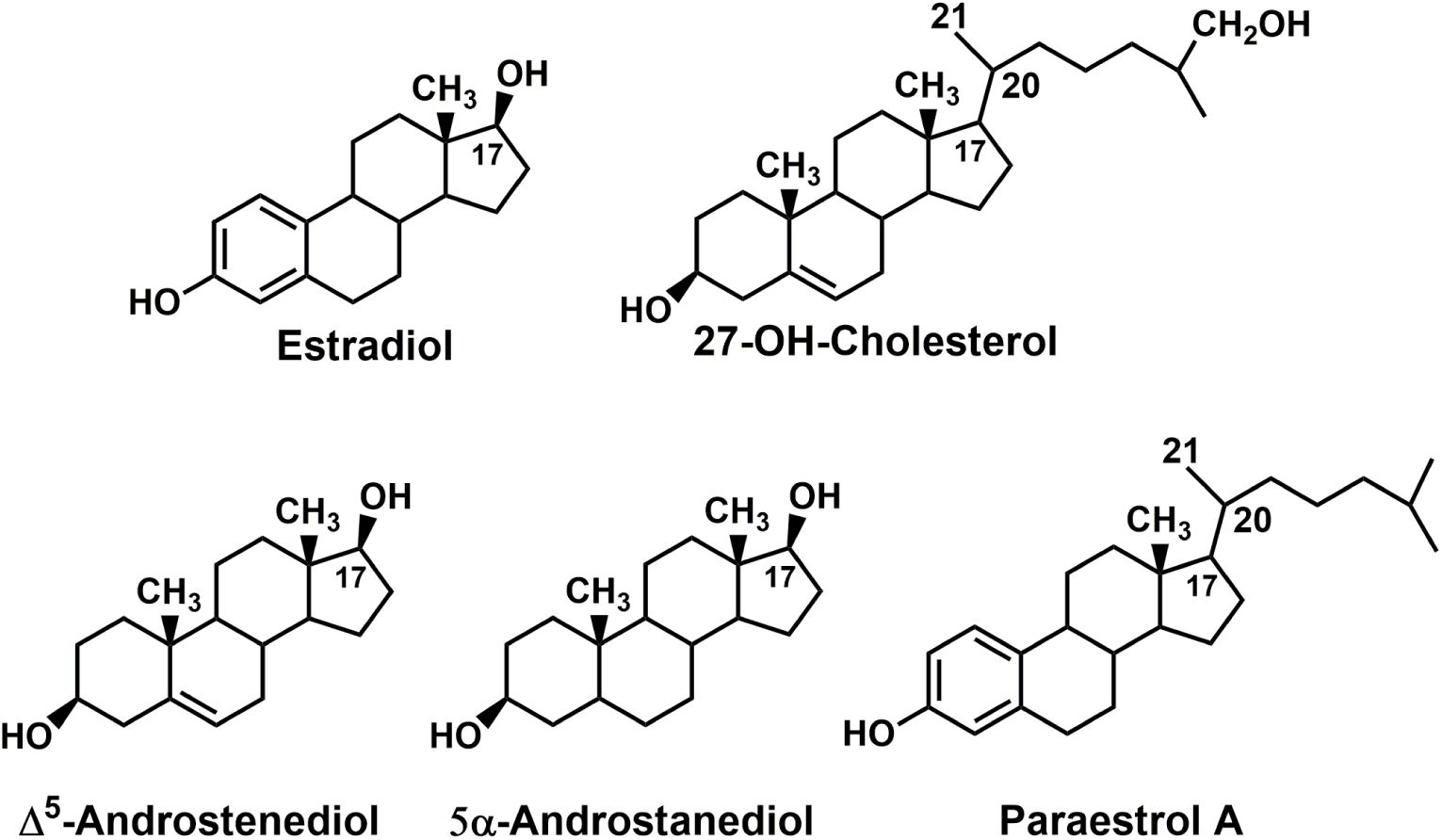
Promiscuity in ligands that are transcriptional activators of the estrogen receptor. The estrogen receptor is activated by steroids with diverse structures [8, 42, 44, 76]. Estradiol, the main physiological estrogen, contains an aromatic A ring with a C3-hydroxyl. By contrast, Δ^5^-androstenediol, 5α-androstanediol, and 27-hydroxycholesterol contain a cyclohexane A ring, with a 3β-hydroxyl and a C19 methyl group. Δ^5^-Androstenediol and 27-hydroxycholesterol contain a B ring with an unsaturated bond between C5 and C6. Δ^5^-Androstenediol, 5α-androstanediol, and 27-hydroxycholesterol are physiological estrogens [27, 46, 50, 61, 67, 80]. Paraestrol A, a cholesterol analog with an aromatic A ring and a C3 phenolic group, has been proposed to be one of a class of cholesterol derivatives that may have been transcriptional activators of the ancestral estrogen receptor [15].

**Figure 2.**
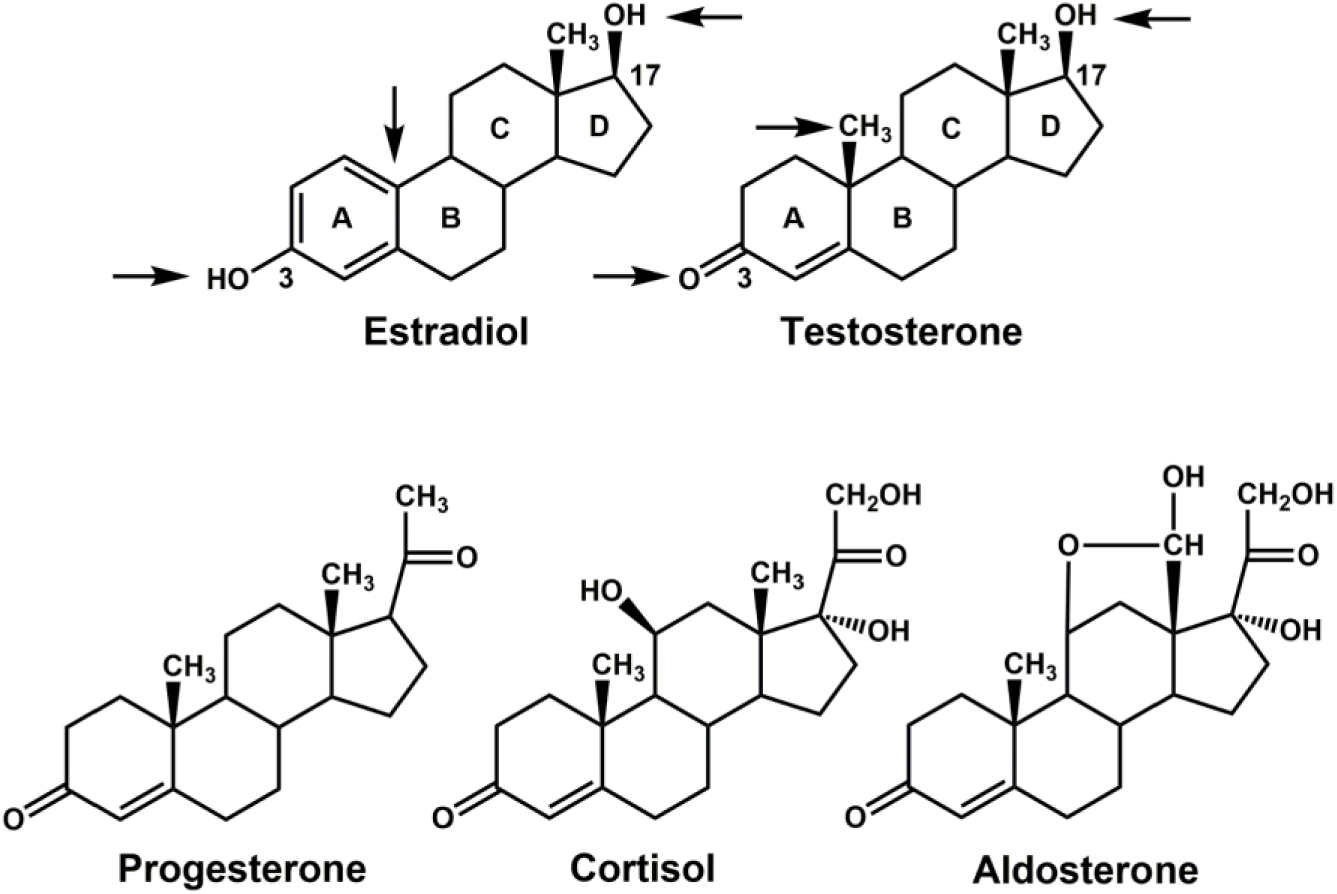
Comparison of estradiol with other sex steroids and adrenal steroids. Testosterone, progesterone, cortisol, and aldosterone contain an A ring with an unsaturated bond between C4 and C5, a C3 ketone, and a C19 methyl group. Estradiol has an aromatic A ring with a C3 hydroxyl and lacks a C19 methyl group. The absence of a C19 methyl group in estradiol allows a close fit between the A ring on estradiol and ERα and ERβ, as seen in the crystal structures of human ERα [34, 36] and ERβ [35, 113].

**Figure 3.**
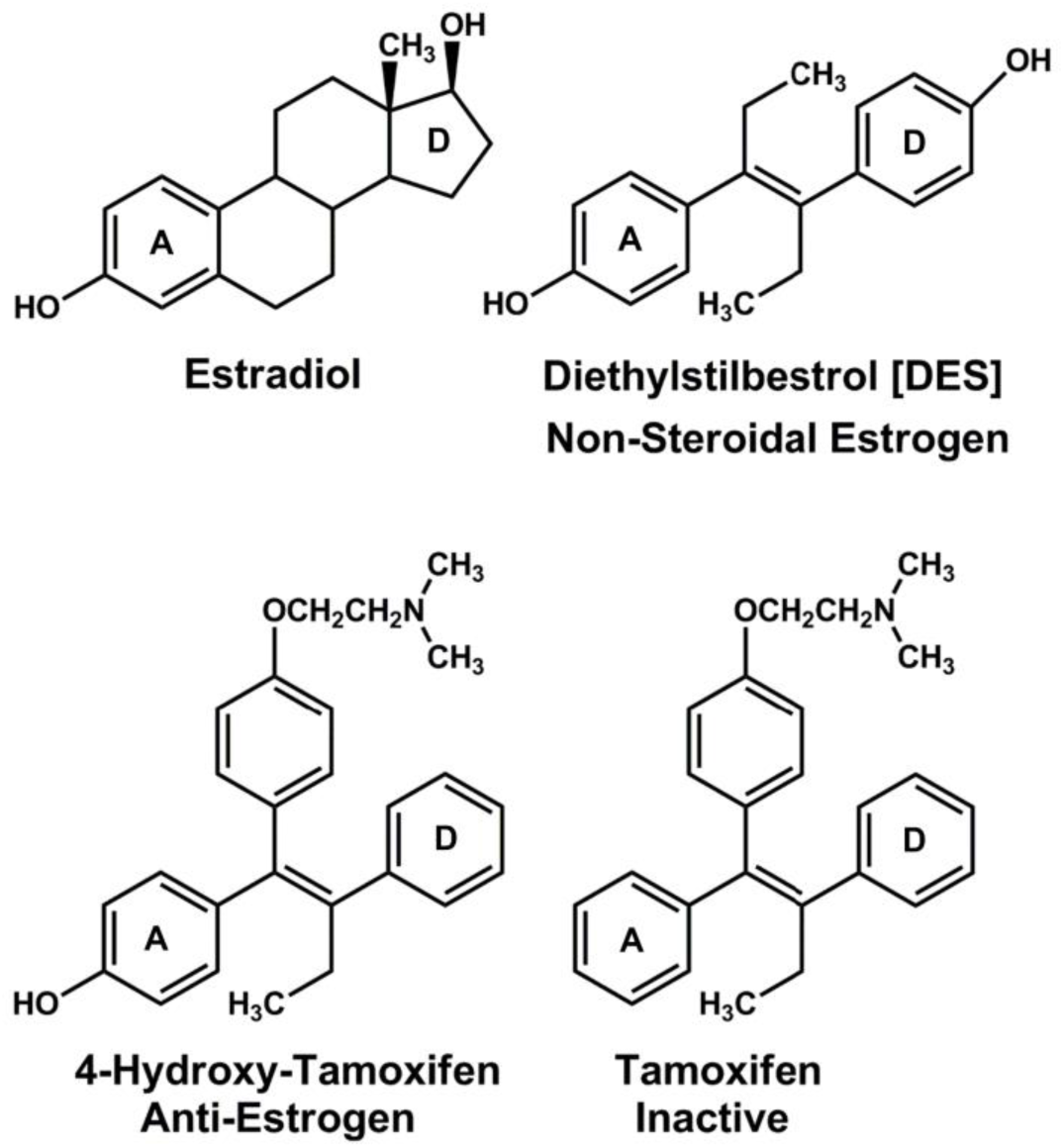
Synthetic estrogens. The role of an aromatic A ring with a C3-alcohol in the biological activity of estradiol provided a template for the synthesis of drugs to regulate estrogen action, such as diethylstilbestrol and 4-hydroxytamoxifen. Diethylstilbestrol is a high-affinity agonist for both ERα and ERβ [27], 4-hydroxytamoxifen is an important anti-estrogen, while tamoxifen, which lacks a phenolic A ring is inactive [27, 42].

Unfortunately, various industrial chemicals containing phenolic groups can disrupt physiological responses to the ER in healthy people, and thus act as endocrine disruptors (Figure 4) [8-12].

**Figure 4.**
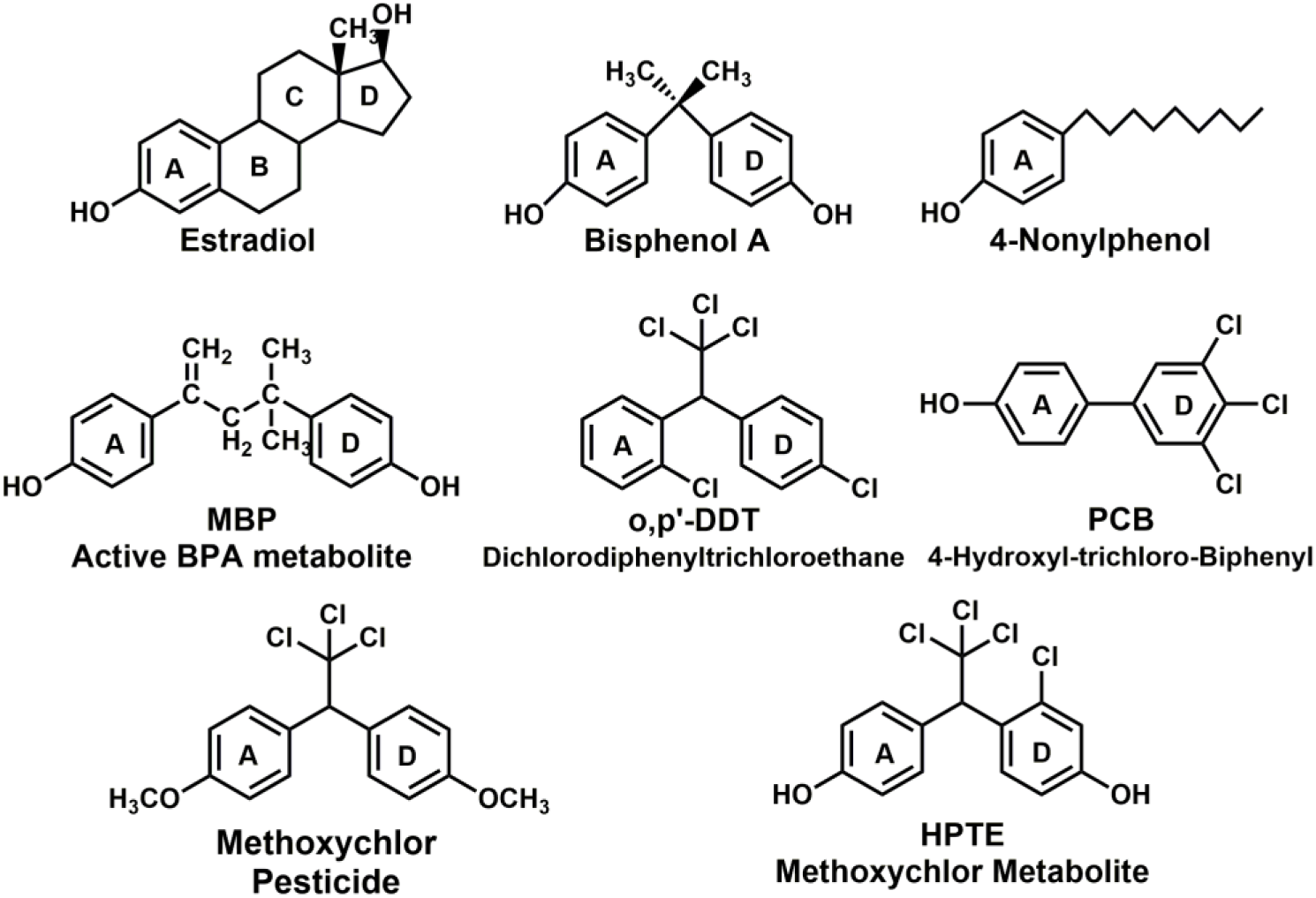
Synthetic chemicals that act as endocrine disruptors. Many synthetic chemicals used in plastics and other industrial products contain a phenolic group that mimic the A ring of estradiol or two phenolic groups that mimic the A and D rings of estradiol [40, 77]. For example, bisphenol A, which is used in plastics and is widespread in the environment, is a xeno-estrogen [40, 41, 114]. BPA can be metabolized to 4-methyl-2,4-bis(4-hydroxyphenyl)pent-1-ene (MBP), which has transcriptional activity at nM concentrations [115, 116]. Metabolism of the pesticide methoxychlor [1,1,1-trichloro-2,2-bis(4-methoxyphenyl)ethane] to HPTE (2,2-bis(*p*-hydroxyphenyl)-1,1,1-trichloroethane) also yields an active xeno-estrogen that is approximately 100-fold more active than methoxychlor [75]. Interestingly, despite the absence of a phenolic group, o,p -DDT (2,2-bis(o,p-dichlorophenyl)-1,1,1-trichloroethane) is a pesticide with estrogenic activity [76, 77].

Interestingly, many natural products in plants also contain phenolic structures, which yield phytochemicals that are transcriptional activators of the ER (Figure 5) [8, 9, 13].

**Figure 5.**
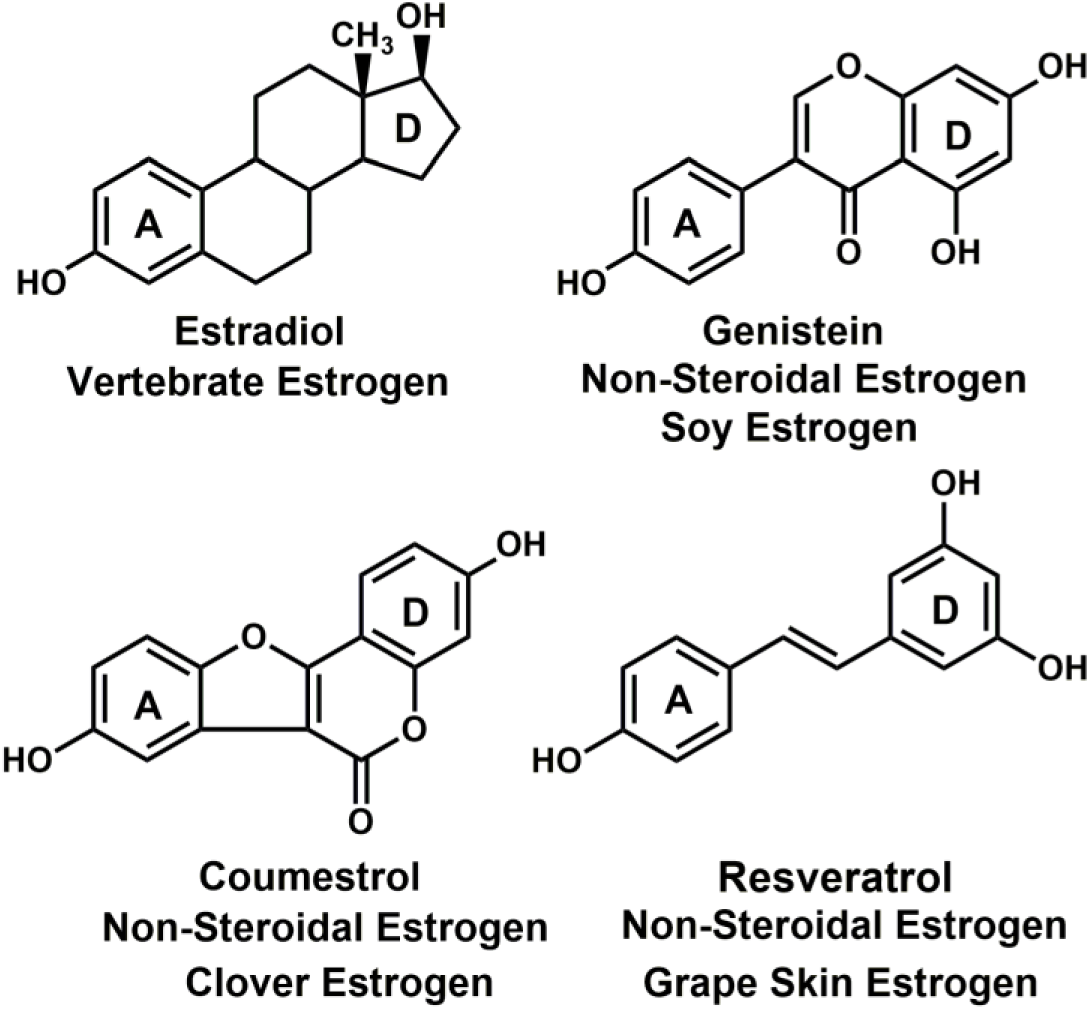
Chemicals from plants that act as estrogens. Many chemicals in plants contain one or two 6-carbon rings with hydroxyl substituents that can mimic estradiol such that the phytochemical is an agonist for ERα and ERβ [8, 40, 42, 43]. Genistein [43] and coumestrol [77] contain aromatic structures that can mimic the A and D rings of estradiol. Resveratrol is a phytoestrogen found in the skins of red grapes [13].

The diversity in the A ring of physiological estrogens (Figure 1) also expands the potential structures of chemicals that can be used as drugs for treating estrogen-dependent diseases in vertebrates, as well as of industrial chemicals that need to be screened for potential disruption of estrogen physiology [9, 14].

Synthesis of 27-hydroxycholesterol and Δ^5^-androstenediol is simpler than that of E2 [4, 9, 15] (Figure 6), and this has important implications for the identity of the estrogen that was the transcriptional activator the ancestral ER [9, 16-18].

**Figure 6.**
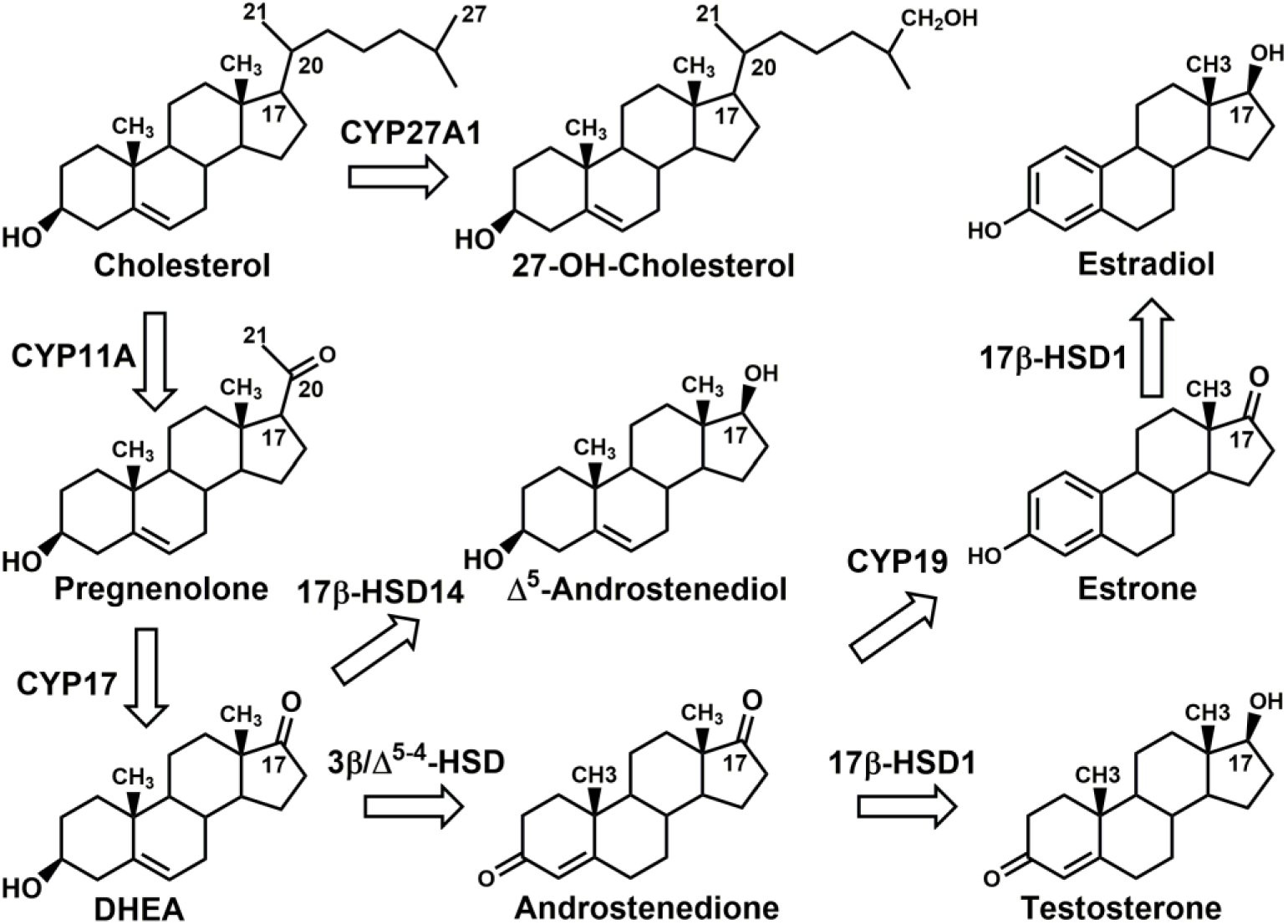
Comparison of pathway for synthesis of estradiol, Δ^5^-androstenediol, and 27-hydroxycholesterol. E2 is synthesized in three enzymatic steps from dehydroepiandrosterone (DHEA) [4, 80]. By contrast, Δ^5^-androstenediol is synthesized from DHEA in one step [46, 53]. 27-Hydroxycholesterol is synthesized in one step from cholesterol.

The earliest ortholog of mammalian ER is found in amphioxus [4, 15, 19-21], a marine chordate that diverged from vertebrates more than 520 million years ago. Based on parsimony, we propose that one or both of these C19-containing steroids or a related metabolite were ligands for the early chordate ER, with E2 evolving later as the canonical estrogen [9, 16]. Many actions of the ER in non-reproductive tissues evolved in vertebrates long before the evolution of mammals, in which reproductive activity of the ER has long been a main focus [22, 23]. An evolutionary perspective of ER signaling [9, 24] provides insights into its promiscuous actions in different tissues: brain, heart, bone, and kidney, as well as traditional ER activities in reproductive organs.

## 2. Estradiol, a hormone with many physiological actions in females and males

Owing to its key role in female reproduction, E2 often is described as a ‘female’ sex hormone. However, E2 also is an important steroid in males [25, 26]. The physiological actions of E2 and other estrogens are mediated by two ER isoforms, ERα and ERβ [27-29]. Through transcriptional activation of these ERs, estrogens regulate variety physiological responses in reproductive organs [22, 23, 26, 28], brain [30-33], bone, and the cardiovascular system [22].

## 3. What is an estrogen?

E2 is the major physiological ligand for the vertebrate ERs [23, 28]. Compared to other adrenal and sex steroids, E2 is a compact hydrophobic molecule with few side chains that could interact with the ER (Figure 2). The aromatic A ring on E2 is unique among adrenal and sex steroids (Figure 2). The crystal structures of ERα and ERβ provide insights into the key contacts between E2 and the ER [8, 34, 35]. Owing to the aromatic chemistry in the A ring, the C3-alcohol can function as a hydrogen-bond donor and acceptor with Arg-394 and Glu-353 in human ERα [8, 34] (Figure 7) and with Arg-346 and Glu-305 in ERβ [35]. In addition, because of the aromatic A ring, the interface between the A and B ring is flat and E2 lacks a C19 methyl group, which allows ERα and ERβ to form a close contact with this part of E2 (Figure 7) [8, 14, 34, 35]. By contrast, testosterone and other 3-ketosteroids (Figure 2) have a different A ring and contain a C19 methyl group, and thus their receptors use a different chemistry to contact the A ring [36-38].

**Figure 7.**
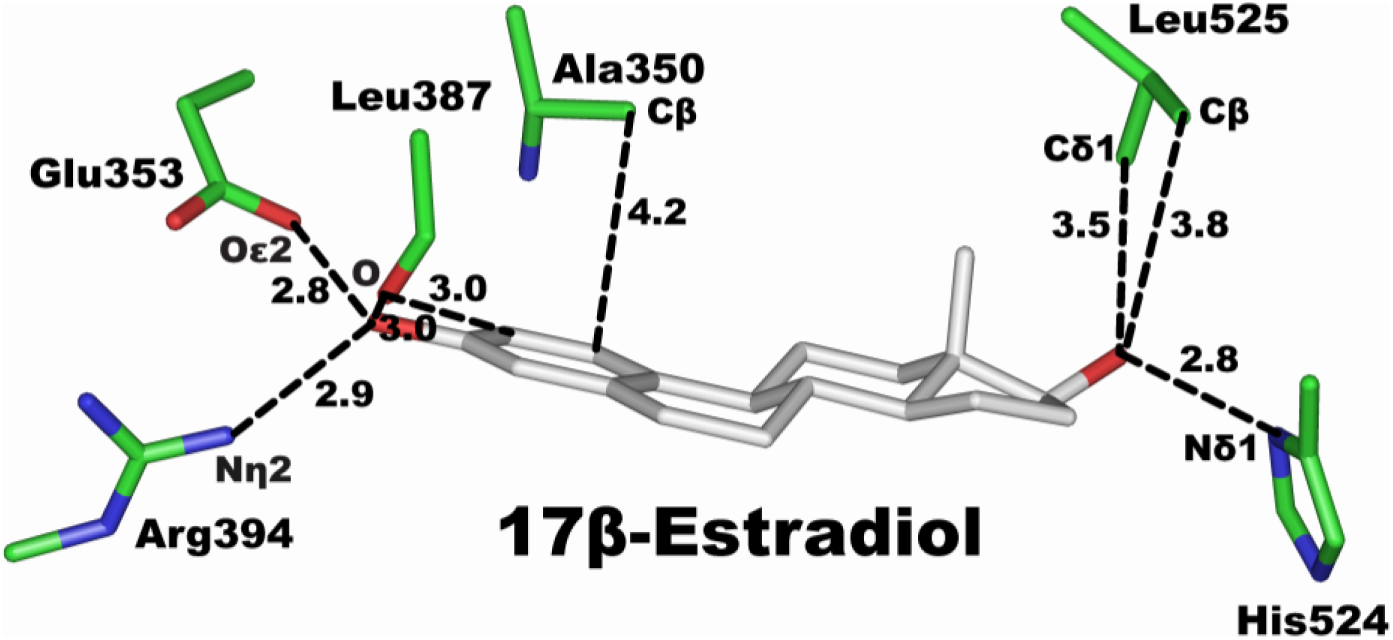
Structure of ERα with 17β-estradiol [34, 117]. ERα (PDB: 1G50) [117] was used for this figure. The C3-hydroxyl on E2 forms hydrogen bonds with Glu-353 and Arg-394 on ERα. The 17β-hydroxyl has a hydrogen bond with His-524.

Also important in binding of E2 to the ER is the hydroxyl on the D ring, which has a stabilizing contact with His-524 in ERα (Figure 7) and with His-475 in ERβ [8, 9, 34, 35].

The spatial relationship of the functional groups on the A and D rings on E2 has been used to develop chemicals containing properly spaced A and D rings that are either estrogens, such as diethylstilbestrol (DES) or anti-estrogens such as 4-OH-tamoxifen (4-OH-TAM) and raloxifene [7, 8, 34, 36, 39] (Figure 3).

Many synthetic chemicals in the environment contain a phenolic group corresponding to the A ring on E2 and often contain a D ring with some similarity to the D ring on E2, which together promotes binding to the ER, disrupting normal estrogen physiology (Figure 4) [10, 11, 40]. For example, the phenolic hydroxyls on bisphenol A (BPA) contact Arg-394 and Glu-353 and His-524 in human ERα (Figure 8) [14, 41].

**Figure 8.**
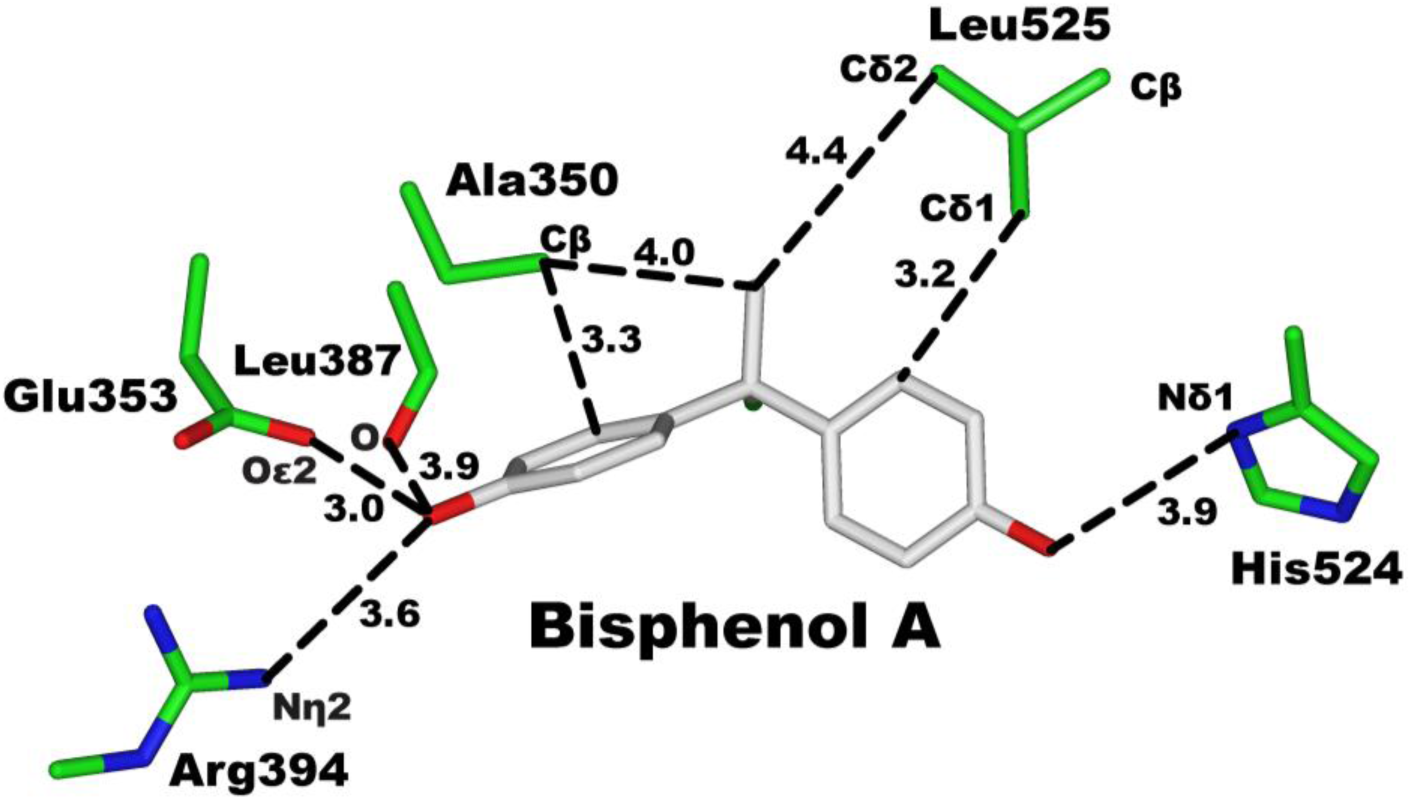
Structure of ERα with bisphenol A. ERα (PDB: 3UU7) [41] was used for this figure. The A ring on BPA forms hydrogen bonds with Glu-353 and Arg-394 in ERα. The hydroxyl on the second ring forms a hydrogen bond with His-524.

Similarly, many chemicals in plants contain an aromatic A ring corresponding to the A ring and a second 6-carbon ring that can mimic the D ring on E2 (Figure 5), which promotes binding with nM K_d_ and transcriptional activation of ERα and ERβ [11, 40, 42]. For example, genistein (Figure 9) [43] and E2 (Figure 7) have similar contacts between their A rings and Arg-394 and Glu-353 in human ERα, and Arg-346 and Glu-305 in ERβ, as well as their D rings and His-524 in ERα, and His-475 in ERβ [14, 43].

**Figure 9.**
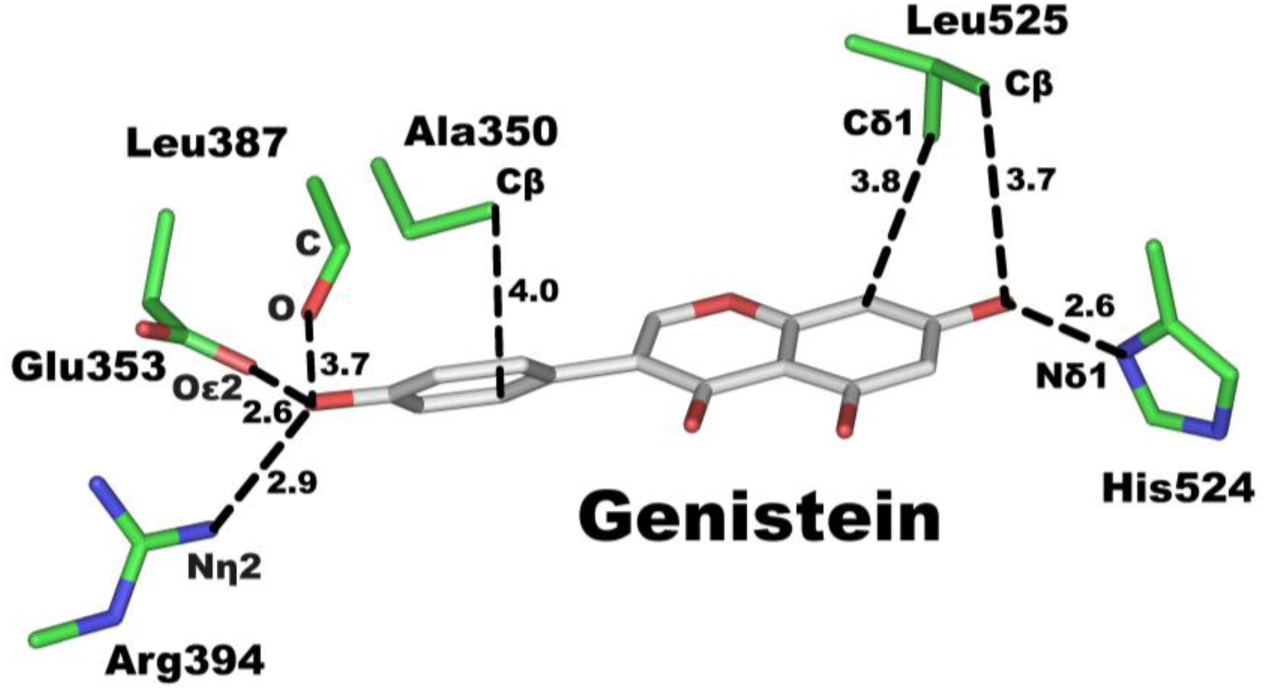
Structure of ERα with genistein. ERα (PDB: 1X7R) [43] was used for this figure. The A ring on genistein forms hydrogen bonds with Glu-353 and Arg-394 in ERα. The hydroxyl group on the D ring on genistein forms a hydrogen bond with His-524.

## 4. Activities of estrogens with novel structures

In the past decade the concept that a steroid must have an aromatic A ring with a C3-phenolic group to be a physiological estrogen has undergone revision because steroids with a different A ring have been shown to be transcriptional activators of ERα and ERβ, and thus act as physiological estrogens [30, 44-47]. Δ^5^-Androstenediol, 5α-androstanediol, and 27-hydroxycholesterol (Figure 1) are examples of physiological estrogens that do not contain an aromatic A ring with a C3 phenolic group, and thus expand the potential structures of physiological ligands for the ER. Phytochemicals and synthetic industrial chemicals also have structures that differ from that of E2. Below we briefly review some physiological activities of these estrogens with novel structures.

### 4.1 Δ^5^-Androstenediol

Δ^5^-Androstenediol (Figure 1) competes with E2 for binding to the ER in rat uterus cytosol [48, 49] and has estrogenic activity in MCF-7 cells [50]. Competitive analysis of binding of Δ^5^-androstenediol to purified human ERα and rat ERβ found that Δ^5^-androstenediol had a K_i_ of 3.6 nM and 0.9 nM, respectively, for human ERα and rat ERβ [27].

Δ^5^-Androstenediol binding to ERβ in astrocytes and microglia inhibits activation of inflammatory response genes [46, 51]. Δ^5^-Androstenediol is synthesized in brain microglia [52, 53]. Inhibition of inflammation in microglia is important in a variety of neurodegenerative diseases [54, 55], including multiple sclerosis, Parkinson’s disease, and amyotrophic lateral sclerosis [56-58]. Interestingly, E2 is not a potent repressor of inflammation in microglia, and 5α-androstanediol has weak activity [46, 51]; that is, Δ^5^-androstenediol has a physiological activity that is absent in E2. Thus, displacement by E2 of Δ^5^-androstenediol from ERβ counteracts the inhibitory effect of Δ^5^-androstenediol on inflammation. This counteracting effect of E2 on Δ^5^-androstenediol inhibition of inflammation in the brain microglia may explain higher incidence of multiple sclerosis in women [51, 59, 60]. Interestingly, binding by the phytoestrogen coumestrol to ERβ (Figure 5) also inhibits inflammation in microglia. However, the phytoestrogen genistein has no effect [46].

### 4.2 5α-Androstanediol

Analysis of binding of 5α-androstanediol to purified human ERα and rat ERβ found that 5α-androstanediol had K_i_ values of 6 nM and 2 nM, respectively, for human ERα and rat ERβ [27]. 5α-Androstanediol binding to ERβ has a role in controlling prostate growth [30, 61, 62] and inflammation in the brain [31, 63]. Indeed, 5α-androstanediol may be a second physiological estrogen in the brain [30, 31, 63].

Owing to the decline in E2 synthesis after menopause, circulating Δ^5^-androstenediol [64-66] may be important physiological estrogens in post-menopausal women.

### 4.3 27-Hydroxycholesterol

27-Hydroxycholesterol appears to be a transcriptional regulator of both ERα and ERβ [47, 67-69]. 27-Hydroxycholesterol binding to ERα reduces bone density [69, 70]. The K_d_ values of 27-hydroxycholesterol for ERα and ERβ are about 1.3 μM and 0.4 μM, respectively [47, 67, 68], which are over 10^3^-fold higher than the K_d_ of E2 for human ERα and ERβ [27]. Levels of circulating 27-hydroxycholesterol are in the range 0.15 to 0.73 μM [47, 67, 71].

27-Hydroxycholesterol acts through activation of ERα to promote the growth of ER-dependent breast tumors [72]. At 10 nM, 27-hydroxycholesterol promoted MCF-7 cell and Ishikawa cell proliferation [73].

### 4.4 Paraestrol A

Paraestrol A is a recently described steroid that resembles cholesterol and with an aromatic A ring (Figure 1). Paraestrol A has been proposed to be an ancestral estrogen [15]. Although this steroid has not been found in mammals, clearly paraestrol A or a metabolite, such as 27-OH-paraestrol A, must be considered as a possible ‘first’ transcription activator of the ER in amphioxus or another chordate in which the first steroid-activated ER evolved [9, 15, 19, 74].

### 4.5 Phytochemicals

Many plants synthesize chemicals that have sufficient structural similarity to E2 so as to bind ERα and ERβ [8, 13, 14, 42] (Figure 5). These chemicals have an A ring with an alcohol at C3 and a 6-carbon D ring. Despite the different D ring structures in genistein (6-carbon ring) and E2 (5-carbon ring), the alcohol on the genistein D ring conserves a key contact with Nδ1 on His524 in ERα (Figure 9). The role of these phytoestrogens in plants is not fully understood, although protection against herbivores through phytoestrogen interference in herbivore physiology is one possibility. There is increased interest in the use of these phytoestrogens in human health [13]; however, their benefits remain controversial.

### 4.6 Environmental chemicals and endocrine disruptors

A major concern regarding the health of the environment is that many synthetic chemicals have structures that allow binding to ERα and ERβ [8, 9, 14, 42] (Figure 4). For example, bisphenol A (BPA) and E2 conserve key binding interactions with ERα (Figure 8). Metabolism of pesticide methoxychlor to HPTE results in an xenoestrogen with approximately 100-fold more activity demonstrating the importance of the phenolic group [75]. Unexpectedly, DDT (dichlorodiphenyltrichloroethane), which lacks alcohol substituents, is nevertheless a xenoestrogen [76, 77]

## 5. An evolutionary perspective. What is the ancestral estrogen and what were its ancestral physiological activities; that is, why is E2 more than a reproductive hormone?

The traditional model is that E2 and its metabolites are the main physiological estrogens in mammals, and thus the estrogenic actions of Δ^5^-androstenediol, 5α-androstanediol, and 27-hydroxycholesterol which lack an aromatic A ring are perplexing. The physiological actions of ERα and ERβ in non-reproductive tissues, such as the brain, also deserve elucidation. To begin to provide a perspective on these phenomena, we draw on Dobzhansky [78] with a modified paradigm ‘Nothing in estrogen physiology makes sense except in the light of evolution’. We use this evolutionary perspective [9, 79, 80] to investigate various inter-related questions including when did the ER arise, what were the ligand(s) that activate the ancestral ER receptor, and what were the physiological actions of the ER in basal chordates and vertebrates?

### 5.1 When did the ER arise?

Nuclear receptors are found in basal animals, but not in plants, yeast, or bacteria [81-86]. Receptors for adrenal and sex steroids evolved in deuterostomes [18, 81, 82, 85-88]. Sequence analysis indicates that the ER evolved before the AR, GR, MR, and PR [4, 18, 74, 87]. Amphioxus, a protochordate in the line leading to vertebrates, contains an ancestral ER and another steroid receptor (SR), which appears to be the ancestor of the 3-ketosteroid receptors [19-21, 89]. Interestingly, E2 does not bind to amphioxus ER. However, E2 is a transcriptional activator of the SR, which is not activated by androgens, glucocorticoids, mineralocorticoids, or progestins [19-21].

### 5.2 What was the ancestral estrogen?

Thus far, only E2 and E1 have been found to be transcriptional activators of amphioxus SR [19, 20]. Other potential estrogens such as Δ^5^-androstenediol, 5α-androstanediol, 27-hydroxycholesterol, and paraestrol A have not been tested for activation of amphioxus SR. The synthesis of these steroids is more parsimonious than the pathway for the synthesis of E2 (Figure 6). In fact, synthesis of testosterone is simpler than that of E2. Indeed, testosterone is an intermediate in the synthesis of E2, suggesting that the AR should have evolved before the ER.

Two hypotheses have been advanced to explain the puzzle that E2 is an ancestral ligand for the SR despite the more complex synthesis of E2 relative to testosterone. Both hypotheses rely on the identity of the ancestral estrogen. In one hypothesis, based on E2 being the ancestral estrogen, Thornton [17-19] proposed the novel ‘ligand exploitation’ model in which E2 was the ancestral ligand for the first ER, and testosterone was inactive because the AR had not yet evolved [19]. Through gene duplication and sequence divergence of an ancestral SR, receptors for the AR and other 3-ketosteroid receptors evolved. In this model, the evolution of the AR ‘exploited’ the presence of testosterone as a biological intermediate in estrogen synthesis [17-19]. Further gene duplications led to the evolution of the other adrenal and sex steroid receptors.

A more conventional model uses parsimony in the pathway for synthesis of estrogens is to identify the ancestral estrogen(s) [9, 16]. The ancestral estrogen is proposed to be Δ^5^-androstenediol, which is synthesized in one step from dehydroepiandrosterone (DHEA) (Figure 6) [9, 16]. Synthesis of Δ^5^-androstenediol does not require either aromatase or 3β,Δ^4–5^-hydroxysteroid dehydrogenase, providing a more parsimonious synthesis than for either E2 or E1, which requires both enzymes.

The evidence that Δ^5^-androstenediol is a high-affinity ligand for the ER supports considering this steroid as a ligand for the ancestral ER. 5α-Androstanediol, 27-hydroxycholesterol, and paraestrol A also may be ligands for the ancestral ER. Of the three steroids with a C19 methyl group, the synthesis of 27-hydroxycholesterol from cholesterol is the most parsimonious [9] (Figure 6). The recent report by Markov et al. [15] of paraestrol A, an aromatized cholesterol derivative, provides another potential ancestral ligand for the ER.

## 6. ERα and ERβ are expression in the brain: transcriptional activation by 5α–androstanediol and Δ5–androstenediol

The presence of an ER and SR in ovaries and testis in amphioxus indicates that some reproductive functions of estrogens in males and females evolved in a basal chordate [19, 20, 90], although transcriptional activation of the ER in uterus or prostate evolved hundreds of millions of years after the evolution of amphioxus, lamprey, and coelacanth. As mentioned above, the ER has physiological actions in the brain. ERα and ERβ are reported to be expressed in different regions in the brain during development in males and females [28, 30, 91-94]. Thus, E2 is a neurosteroid, as well as a reproductive steroid. Moreover, Δ^5^-androstenediol and 5α-androstanediol, which have a different A ring than E2, have been found to be transcriptional activators of the brain ER [30, 46].

The estrogenic activity of steroids with a C19 methyl group may provide a selective advantage in estrogen physiology that is not provided by E2. In this regard, 5α-androstanediol has been proposed to be a second physiological estrogen in fetal mouse brain based on the timing of ERβ synthesis in mouse brain, which begins at embryonic day 10.5 [30, 95, 96], well before embryonic day 18.5 when aromatase (which catalyzes the synthesis of E2) is first expressed. Thus, there are at least two physiological estrogens in mouse brain. Moreover, other studies find that ERβ mediates the actions of 5α-androstanediol in human brain cells [31, 63].

In addition, Δ^5^-androstenediol is synthesized in the brain [52, 53], providing a third endogenous estrogen in mouse brain. Finally, 27-hydroxycholesterol also is present in the brain [97]. Thus, there may be four physiological neuro-estrogens. The unique anti-inflammatory response in brain microglia due to binding of Δ^5^-androstenediol to ERβ may be the first of a series of specialized physiological activities for Δ^5^-androstenediol, 5α-androstanediol, and 27-hydroxycholesterol in the brain and other organs. It is interesting that E2 and Δ^5^-androstenediol regulate different physiological responses through ERβ in microglia.

We propose that some activities of the ER in different regions of the mammalian brain [30, 32, 33, 98] were important in amphioxus or in lamprey. Although the head and brain are poorly developed in amphioxus [99-102], many genes in amphioxus are expressed in the brain in lamprey and other vertebrates [100, 102, 103]. It appears that these genes were important in the evolution of the brain during the transition from amphioxus to lamprey [100-102]. It is tempting to speculate that transcriptional regulation of the SR in an ancestral amphioxus by Δ^5^-androstenediol, 5α-androstanediol, 27-hydroxycholesterol, or paraestrol A, as well as by E2, was important in brain evolution [79, 80], which would place the origins of responses to estrogens in the brain as contemporary with its reproductive actions.

Also relevant for the evolution of estrogen physiology is the presence in lamprey of a PR and corticoid receptor (CR) [4, 18, 104], which is the ancestor of the GR and MR [4, 104]. These steroid receptors have transcriptional properties that similar to their mammalian orthologs [38, 105]. Crosstalk between the ER and the PR, GR, and MR may have contributed to the evolution of the brain [106-108] and other organs during the transition from amphioxus to lamprey.

## 7. Future research

There is a need to verify the presence of Δ^5^-androstenediol, 5α-androstanediol, and 27-hydroxycholesterol in amphioxus and lamprey to provide evidence for the physiological activity of these 3-ketoestrogens early in the evolution of the ER. Synthesis of Δ^5^-androstenediol, 5α-androstanediol, and 27-hydroxycholesterol in amphioxus and lamprey can be studied through metabolism of radioactive cholesterol in tissue extracts from amphioxus and lamprey. Of course, transcriptional activation of amphioxus SR and lamprey ER by Δ^5^-androstenediol, 5α-androstanediol, and 27-hydroxycholesterol will be necessary to determine their biological activity in relation to E2. The functions of 3-keto-estrogens in mammals merit further study in view of the evidence that 5α-androstanediol and Δ^5^-androstenediol are important in prostate [61] and brain microglia [46, 51], respectively. Investigation of Δ^5^-androstenediol, 5α-androstanediol, and 27-hydroxycholesterol actions in tissues with ERα and ERβ during different stages in development may uncover additional physiological responses mediated by these ‘alternative’ estrogens. In this regard, the slow continuous increase in DHEA synthesis from birth, which is coupled with increased synthesis of Δ^5^-androstenediol [109], merits further study, as does the role of Δ^5^-androstenediol during fetal development [110], when DHEA levels are high [111, 112].

## Author contributions

M.E.B and R.L. wrote and edited the MS.

## Funding

M.E.B. was supported by Research Fund #3096.

## Competing interests

We have no competing interests.

